# Not a one-way road – severity, progression and prevention of firework fears in dogs

**DOI:** 10.1101/654301

**Authors:** Stefanie Riemer

## Abstract

Noise fears represent a highly prevalent welfare problem in dogs. An online survey was performed to explore severity and progression of firework fears in dogs, and relationships with demographics, health, behaviour problems, and owners’ training efforts to prevent or alleviate firework fears. 1225 responses were analysed. Fifty-two percent of dogs were at least partially affected by firework fears, and the great majority developed a fear of fireworks in the first year of life, with a decreasing frequency of new occurrences up until seven years, and only few newly affected dogs beyond this age. While almost three quarters of fearful dogs had recovered by the next morning, recovery took up to one day in 10%, up to one week in 12%, and several weeks or even months in >3%. Univariate analyses indicated a significant effect of breed group, age, sex, neuter status, origin and age at acquisition on severity of firework fears in dogs. However, binomial models including multiple predictors of presence/ absence of firework fears identified only age, breed group (mixed breeds being most affected), health problems, and an interaction between health problems and age as significant predictors. This discrepancy might be explained by collinearities of predictors and underlying differences between mixed-breed dogs and purebreds, such as mixed breeds being acquired from shelters more often, being adopted at higher ages, and being neutered more often. Firework fears are highly correlated with fears of gunshots and thunder, and to a low extent with fears of other noises, but not with any other behavioural problems. Both improvement and deterioration of firework fears were frequently reported. While an early age of onset and breed differences point to a strong genetic contribution to firework fears, training puppies or non-fearful adults to associate the noise with positive stimuli is highly effective in preventing later development of firework fears.

## Introduction

Fear of noises is highly prevalent in dogs and represents a significant welfare concern, with up to half of the pet dog population affected (1–3). Yet, only a minority of pet owners seem to seek professional advice regarding this issue (2,4), and while some studies investigated treatment options (e.g. (5–10), there is a lack of research on preventive measures.

At present, no unified terminology exists in the field and different authors refer to fear of noises as “noise sensitivity” (e.g. (11), “noise reactivity” (e.g. (12), “noise aversion” (13) or “noise stress” (reviewed by (12). Further distinctions are often made between “fear” (an adaptive response to a stimulus considered to be potentially dangerous), “anxiety” (anticipation of a negative outcome, lacking a specific eliciting stimulus)(1,14), and “phobia” (an extreme, long-lasting reaction which can be elicited by a low stimulus intensity and in human psychiatry is considered irrational (reviewed by (2)). However, as Overall et al. (12) point out, studies are typically lacking sufficient criteria to differentiate between terms such as “reactivity” or “phobia” in dogs, and moreover even in the most commonly used rodent model species to study fear and anxiety, the states of fear and anxiety can often not be differentiated behaviourally (reviewed by (14)). Therefore, in the remainder of the paper I will use the term “noise fears” or “firework fears” to denote any fearful, anxious, stressed or phobic reactions of dogs when exposed to noises or fireworks, respectively.

Fireworks appear to be the most common trigger of noise fears in dogs, although the great majority of affected dogs concomitantly show fears of gunshots and thunderstorms (1,2,12). Breed or breed group has consistently been identified as being associated with different susceptibility to firework fears (1–3,12), pointing to contributing genetic factors. While the observed breed differences in noise fears are likely polygenic (12), a few genes contributing to noise fears in some breeds have recently been identified (15,16). Meanwhile, the fact that crossbreeds were the group with the highest incidence of firework fears in (2) points to possible environmental influences (i.e. socialisation experiences) associated with the dogs’ origin.

Besides breed, age is a common risk factor, with the prevalence of firework fears increasing with age (1–3). While fear of noises is usually observable at an early age (median onset: 2 years in (3)), one possible factor contributing to the development of noise fears may be pain, and this may explain why in some dogs an onset of noise fears occurs at a later age: In a recent study comparing ten noise-sensitive dogs with an underlying muscoskeletal pain problem and ten noise-sensitive dogs with no detectable pain, the age of onset was almost four years later in the painful dogs (mean 6.5 years), compared to the non-painful dogs (mean 2.67 years)(17).

Regarding the effects of sex and neutering on noise fears in dogs, studies have yielded inconsistent results. Both intact females and neutered dogs of both sexes had a higher incidence of noise fears in (1); in contrast no effect of sex or neutering was found in other studies (2,4). Origin of the dog (e.g. rescue, pet shop, breeder etc.) could be expected to affect fearfulness in dogs due to likely being associated with differential socialisation experiences; nonetheless, this variable did not influence degree of firework fears in (4). In (2), the only significant effect of origin was for dogs bred by their current owner, which had a lower incidence of noise fears compared to dogs from other sources (breeder, rescue centre, or other including pet shop).

Although a number of publications indicate behavioural signs shown by dogs when exposed to loud noises (e.g. trembling, freezing, panting, salivation, lowered body posture, tucked tail, hiding, escape attempts, social withdrawal, pacing, involuntary elimination, and destructive behaviour with or without self-injury (3,12,13,18–21), there is a lack of knowledge on how long such behavioural changes as a result of firework exposure typically persist. One study reports that the median duration of behavioural changes in the aftermath of a firework was two hours (with a mean of 1.83, SD 0.044, (4)), but little is known on the distribution of responses, and how many dogs may be affected beyond the timeframe of a few hours. This detail is, however, of great importance in relation to dogs’ welfare. Whenbehavioural effects persist beyond the time of direct exposure, especially if they last several days or even weeks to months, this would indicate a significant welfare impairment.

Another question of interest relates to the progression of firework fears in dogs, i.e. are they typically stable once developed or is a deterioration inevitable? It has been shown that behaviour problems other than noise fears, including fearfulness (encompassing fear towards people, dogs, handling and non-social fear) and aggression, typically increase over time (22). Regarding noise fears, some small-scale studies investigated the effects of therapeutic interventions (e.g. (6,10,21,23,24), but only one larger scale questionnaire study asked owners to describe dogs’ changes in firework fears over time, and this study used change in fear as dependent variable in further analyses without providing descriptive statistics (4). The survey by Blackwell et al. (2) indicated that spontaneous recovery from noise fears may be possible in a small number of individuals, although for half of these animals, loss of hearing appeared to be the responsible factor. Thus, although it is generally assumed that noise fears usually get worse over time (e.g. (12), in line with the finding that noise fears increase with age (1–3), very little is known about patterns of progression at a population level.

Finally, despite a number of publications on intervention to treat firework fears in dogs (5–10), an important issue that seems to be comparatively neglected is how to prevent fears of fireworks in dogs from developing in the first place. One study indicated that playing the radio during feeding time in German shepherd puppies between the ages of 16 and 32 days led to more favourable responses to sudden loud noises when tested in a puppy test at the age of seven weeks (25). On the other hand, no beneficial effects of gradually increased auditory stimulation from the age of three weeks was found when 7-week old puppies (German shepherds, Belgian Malinois, Dutch shepherd, and crosses between these breeds) were exposed to sudden noises in a behaviour test (26).

For working dogs, it has been recommended to gradually introduce potentially fear-provoking stimuli to achieve habituation, while pointing out that “the relative risk of habituation or sensitization will vary with characteristics of the stimulus, the personality of the dog, and the state of the individual animal at the time of stimulus presentation” (27). Initially, the stimulus intensity should be chosen as low as possible to ensure that it is below all animals’ startle threshold. Animals with already established fears should be identified through preliminary tests and undergo tailored training “including desensitization, counterconditioning, prior to controlled exposure, and habituation” (27). However, more research is needed on how to create the most resilient dogs, both when puppies are still at the breeders, as well as when they are with their new owners or handlers.

Thus, the current study aimed to investigate

1. prevalence and severity of firework fears in pet dogs, age of onset and demographic influencing factors, as well as time until recovery after firework events
2. co-occurrence of firework fears with other behavioural problems and
3. progression and prevention of firework fears in pet dogs, and whether this could be influenced by owners’ training efforts.

## Methods

### Ethics statement

Participation in the questionnaire study was voluntary and participants were informed that they could quit the survey any time, and that no data would be saved until they hit the “Submit” button at the end. Respondents were not required to disclose any personal information other than the country of origin and their level of experience with dogs. All respondents whose answers were included in the analyses gave their consent for the information provided to be used for scientific analysis. For these reasons, no ethical approval was required for the study.

### Questionnaire survey

An online questionnaire survey (in an English and a German version) was distributed to a sample of dog owners via our research group’s website and social media. The advertisement stressed that dogs both with and without fear of fireworks were of interest, aiming to avoid a response bias towards owners of dogs that were affected by firework fears.

The questions covered the owners’ consent for the use of their data, demographic data about the dogs (date of birth, sex, neuter status, breed, country, source of dog, age at acquisition) and dogs’ health problems. For breed groups, the FCI classification was used. If at least ten individuals belonged to a FCI group, they were subsumed as that group; otherwise they were categorised as “Other”. In mixed breeds, dogs with parents from the same FCI group were categorised as belonging to that FCI group, while crosses of parents from different FCI groups or of unknown breed origin were grouped as “mixed breeds”. For the FCI Group 2 (Schnauzer and Pinscher), there were enough individuals from the sections “Pinscher” (N=15) and “Molossians” (N=52), respectively, to warrant including these sections as separate breed groups, and likewise I differentiated between “Retrievers” (N=103) and “Flushing dogs” (N=20) of the FCI Group 8 “Retrievers – Flushing Dogs – Water Dogs”.

To gather information on potential behavioural problems, owners were asked to rate their level of agreement with a number of statements (for example “My dog is afraid of other dogs” or “My dog defends resources against humans”) on a 5-point Likert scale ranging from “disagree strongly”, “tend to disagree”, “partly/partly”, “tend to agree” to “agree strongly”.

#### Firework fears: Welfare impaired score

One main dependent variable for the analyses is the “Welfare impaired score”, which is based on the question “Please rate your level of agreement with the following statement: The overall welfare of my dog is strongly compromised by fireworks”. As the abovementioned scores, it was answered on a five-point Likert scale from “disagree strongly” to “agree strongly”.

#### Firework fears: Fear progression score

The other main dependent variable for analysis is the “Fear progression” score, which was based on the question “How has your dog’s fear of fireworks progressed in the last years?”, with the following response options “My dog was never afraid of fireworks”, “The fear has improved greatly”, “The fear tends to have improved”, “The fear has remained the same”, “The fear tends to have become worse”, “The fear has become much worse” or “I don’t know”. In the subsequent analysis, this question was only analysed for dogs that were affected by firework fears (Welfare Impaired score ≥ 3), so the answers “My dog was never afraid of fireworks”, and “I don’t know” were removed from the sample.

Further questions relating to firework fears included “How long does it take until your dog’s behaviour is completely back to normal following a fireworks?” and “At what age did fear of fireworks first become apparent in your dog?” Owners were also asked whether they had attempted any training to prevent or treat firework fears in their dogs, and if so whether training was commenced when the dog was still a puppy, an adult (before the onset of any firework fears), or after the dog had already shown a fear of fireworks. All relevant questions are available in S1 Table.

### Analysis

Statistica 6.1 (Statsoft Inc. 1984–2004) was used to calculate non-parametric statistical tests, IBM SPSS Statistics Version 23 (© IBM Corporation and its licensors 1989, 2015) was used to calculate Principal Components Analyses (PCA), and R version 3.3.3 (© 2017 The R Foundation for Statistical Computing) was used to compute binomial models.

For the purpose of analysis, the Likert responses were converted to numbers of 1 (“disagree strongly”/ “The fear has become much worse”) to 5 (“agree strongly”/ “The fear has improved greatly”). Health problems received a binary score (0 – no health problems; 1 – one or several health problems).

#### Age of onset of firework fears

To calculate the relative frequency of age of onset of firework fears in dogs, for each age group I calculated the proportion of dogs for whom the onset of firework fears was reported at that age, divided by the total number of dogs having reached this age in the sample.

#### Demographic influencing factors

In order to assess associations between demographic and training factors with severity or progression of firework fears in dogs, non-parametric statistics were performed as the dependent variables were ordinal scores, but not all model assumptions for ordinal models were met. Kruskal Wallis tests were used to test for differences between dogs of different sexes/ neuter status (four groups: male intact, male neutered, female intact and female neutered dogs), differences between breed groups, and between dogs of different origins (e.g. homebred, large-scale breeders, rescue abroad etc.; see details in S1 Table).

However, the non-parametric approach allowed to test the effects of only one factor at a time. Yet it cannot be ruled out that some of the predictor variables, such as source of dogs, breed and neuter status, might be confounded, with dogs obtained from rescue shelters being more likely to be neutered and of mixed breeds. Furthermore, an interaction between age and health problems might be expected in view of a recent study suggesting that onset of noise fears occurred at higher ages in dogs with muscoskeletal pain compared to those not affected by muscoskeletal pain (17).

Therefore, to address the possibility of interactions between some predictors, selected binomial logistic regressions (function glm in R) were calculated with the predictors Sex*Neuter status, Age*Health problems, breed group, source of dog, and age at acquisition as independent variables and a binary “Welfare Impaired” score (re-classified from the 5-point scale) as dependent variable. For this binary score, dogs with a Welfare Impaired score of 1-2 were considered as “not fearful” (0), while dogs with a Welfare Impaired score of 3-5 were considered as “fearful” (1). A step-wise model selection approach based on Akaike’s information criterion (AIC) was used to select the best model.

#### Relationship of firework fears with other behavioural problems

A principal components analysis (PCA) was performed over all questions relating to behavioural problems other than fireworks. As the main aim here was not data reduction, but to detect structure in the data, the number of components retained was not based on a Scree plot or Eigenvalues, but a higher number of components that were biologically meaningful were retained. These seven components covered 83.5% of the variance in the data. Spearman rank correlation tests were performed to assess correlations of the components with the Welfare Impaired score.

#### Effect of training

The effect of training was firstly assessed for the full sample (Welfare Impaired scores ranging from 1-5). The Welfare Impaired score was compared for dogs having received targeted training to prevent noise fears as puppies, those having received such training as adults before developing any noise fears and dogs that did not receive preventative training, using a Kruskal Wallis test. Secondly, in dogs already affected by firework fears (Welfare Impaired scores of 3 and above), a Mann Whitney U test was conducted to compare Fear Progression scores in dogs that had received behavioural training compared to those that had not.

#### Correction for multiple testing

Even when applying the conservative Bonferroni correction for multiple testing, all significant results remained significant, and the original p-values are reported in the Results. For Kruskal Wallis tests, post-hoc testing for between-group differences was performed using Statistica’s inbuilt algorithm after (28), and adjusted p-values are reported.

## Results

### Descriptive statistics

After removing dogs younger than one year at the time of the questionnaire response, 1225 valid responses were obtained, including 527 English and 699 German responses. Subjects included 588 females (of which 430 neutered and one of unknown neuter status) and 637 males (of which 424 neutered and 6 chemically castrated). Dogs that were chemically castrated were not considered for analysing the effect of neutering as we cannot be sure whether they were hormonally equivalent to surgically neutered dogs.

The dogs were of various breeds or mixes, with 729 belonging to a single breed group and 485 being mixed breeds or crosses from parents of different breed groups. Among dogs from a single breed group 61.0% were neutered, while the proportion of neutered dogs in the mixed breeds was higher at 83.5%. Purebred and mixed-breed dogs also differed in the proportions coming from different sources, as shown in S2 and S3 Tables. In particular, mixed-breed dogs were more likely to originate from rescues either locally or abroad and were more likely to be former street dogs, while purebreds were more likely to originate from a breeder (big/ small), from a private person whose bitch had a litter, to have been rehomed from a private person, or to have been bred and raised by their current owner.

### Prevalence of firework fears and age of onset

Based on a Welfare Impaired score of 3 or higher, 639 dogs of the 1225 dogs in the sample (52.2%) were considered to be fearful of fireworks (Table 1).

**Table 1.** Distribution of Welfare Impaired score in the population (based on the statement “The overall welfare of my dog is strongly compromised by fireworks”: 1= strongly disagree; 5=strongly agree)

| **Welfare Impaired**<br>**Score** | **N** | **% of dogs** |
| --- | --- | --- |
| 1 | 385 | 31.43 |
| 2 | 201 | 16.41 |
| 3 | 115 | 9.39 |
| 4 | 140 | 11.43 |
| 5 | 384 | 31.34 |
| **Total** | 1225 | 100 |

148 of the fearful dogs had been adopted as adults when they already showed a fear of fireworks so the age of onset was unknown. In the remaining “fearful” dogs with both current age and age of onset recorded (N=395), there was a clear trend showing that firework fears tend to develop already at a young age: 45% of owners reported that their dogs developed a fear of fireworks already under one year of age. The second most common age of onset was at two years, followed, by three years and four to six years (Table 2). Above six years, very few dogs showed first signs of firework fears. Accordingly, the median age of onset was one year. Table 2 shows the number of dogs in the sample having reached the respective ages at the time of the questionnaire and the numbers and proportions of dogs having experienced an onset of firework fears at the different ages.

**Table 2.** Frequency of onset of firework fears in dogs at different ages, relative to the number of dogs having reached the respective ages in the sample (non-fearful dogs and dogs with missing data for age or age of onset are excluded from the dataset).

| Age | <1 | 1 | 2 | 3 | 4 | 5 | 6 | 7 | 8 | 9 | 10 | 11 | 12 |
| --- | --- | --- | --- | --- | --- | --- | --- | --- | --- | --- | --- | --- | --- |
| Number of dogs currently at this age in the sample | - | 15 | 32 | 46 | 41 | 40 | 51 | 33 | 46 | 28 | 26 | 17 | 11 |
| Number of fearful dogs having reached this age | 395 | 395 | 380 | 348 | 302 | 261 | 221 | 170 | 137 | 91 | 63 | 37 | 20 |
| Number of dogs first developing fear at this age | 178 | 109 | 84 | 46 | 25 | 22 | 16 | 2 | 2 | 1 | 2 | 1 | 1 |
| Proportion of fear started in dogs having reached this age | 0.45 | 0.28 | 0.22 | 0.13 | 0.08 | 0.08 | 0.07 | 0.01 | 0.02 | 0.01 | 0.03 | 0.03 | 0.05 |

### Time until recovery

11.9% of the fearful dogs were reported to behave normally immediately after firework exposure, with 21.6% taking up to half an hour to recover and 17.5% taking up to an hour. 10.3% and 12.6%, respectively, recovered within three hours or by the next morning. The dogs’ behaviour normalised in the course of the next day in 10.4% and in up to three days also in 10.4%. 1.8% were affected for up to one week, 2.3% for several weeks, and 1.2% even for several months, with the latter group including one dog whose behaviour never normalised according to the owners.

### Demographic influencing factors

A Kruskal Wallis ANVOA comparing male intact, male neutered, female intact and female neutered dogs was highly significant (N=1218, H=28.89, p<0.0001). Post-hoc individual comparisons (with adjusted p-values for multiple testing) demonstrated significant differences between male intact and male neutered animals (z=3.208, p=0.008), as well as between female intact and female neutered animals (z=3.677, p=0.001). However, there were no differences between either intact individuals of both sexes (z=0.043, p=1.0) or neutered individuals of both sexes (z=0.968, p=1.0). To summarise, the Welfare Impaired score was significantly higher in neutered dogs of both sexes, but did not show a difference between male and female animals.

The Welfare Impaired score was significantly positively correlated with age, although the strength of the correlation was weak (Spearman Rho=0.20, N=1095, p<0.000001). There was also a significant correlation of the Welfare Impaired score with the dog’s age at acquisition, albeit with an even lower correlation coefficient (N=1225, Rho=0.119, p=0.000026). The Welfare Impaired score differed significantly between breed groups (Kruskal Wallis, N=1220, H=66.163, p<0.0001; Fig 1). Post-hoc tests (with p-values corrected for multiple testing) indicated that mixed breeds had the highest average Welfare Impaired scores, which differed significantly from companion dogs (z=3.531, p=0.038), molossians (z=4.493, p=0.0006), retrievers (z=5.012, p=0.00005) and hounds (z=3.622, p=0.027). Also herding dogs ranked significantly higher on the Welfare Impaired score than molossians (z=3.489, p=0.041) and retrievers (z=3.587, p=0.030) (S4 and S5 Tables).

**Fig 1.**
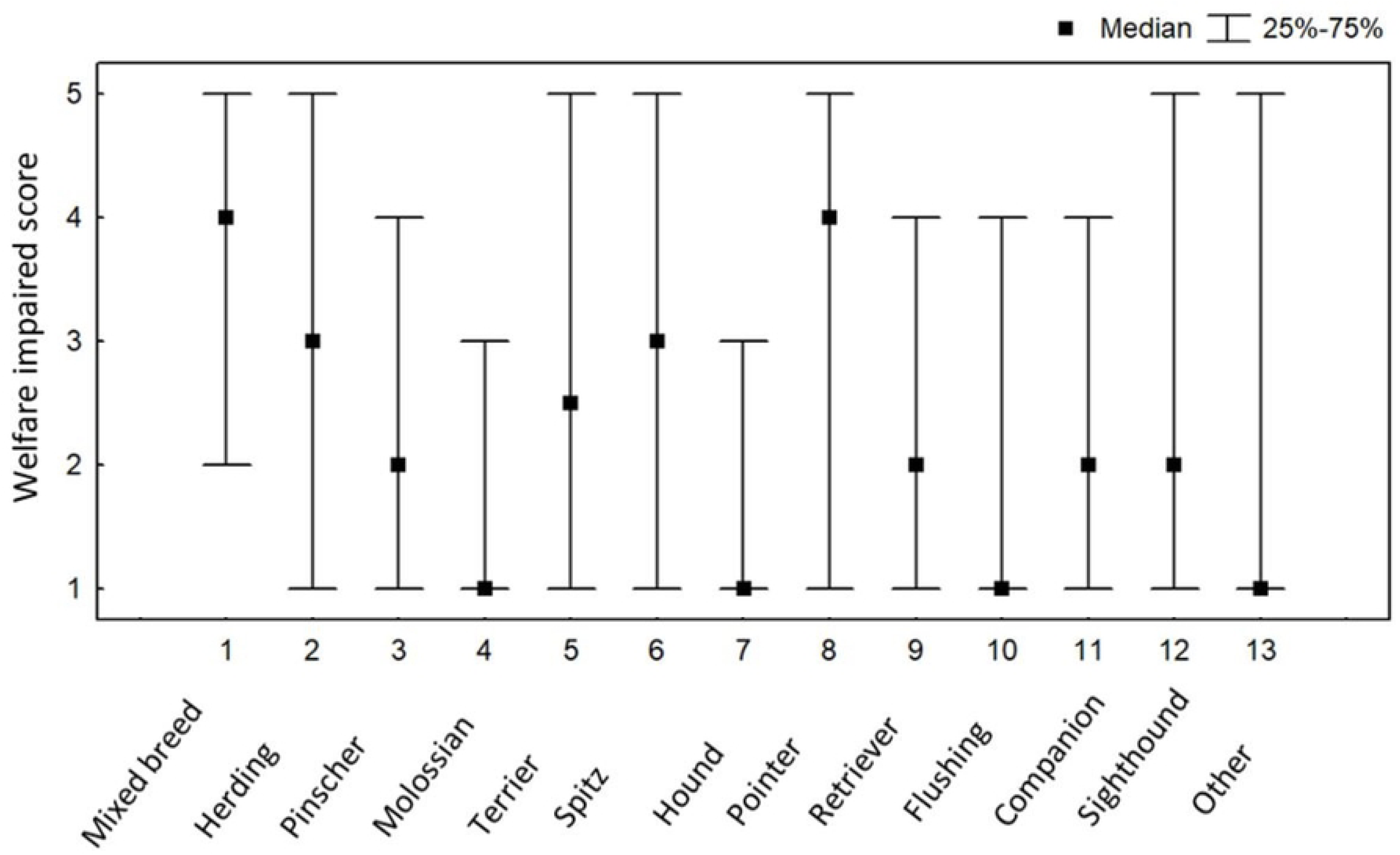
Median Welfare impaired scores and interquartile ranges for the different breed groups in the sample.

There was a significant effect of the source of the dog on the Welfare Impaired score (Kruskal Wallis H=31.715, N=1042, p=0.0001; Fig 2). Dogs that were homebred and retained by their breeders scored the lowest on Welfare Impaired during fireworks. Dogs obtained as adults from rescue organisations or shelters, both within the home country and abroad, had the highest scores, and post-hoc tests (with adjusted p-values to correct for multiple comparisons) indicated that this difference was significant in comparison to dogs from small-scale breeders (local rescues: z=3.483, p=0.018; rescues abroad: z=4.207, p=0.000931)(S6 and S7 Tables).

**Fig 2.**
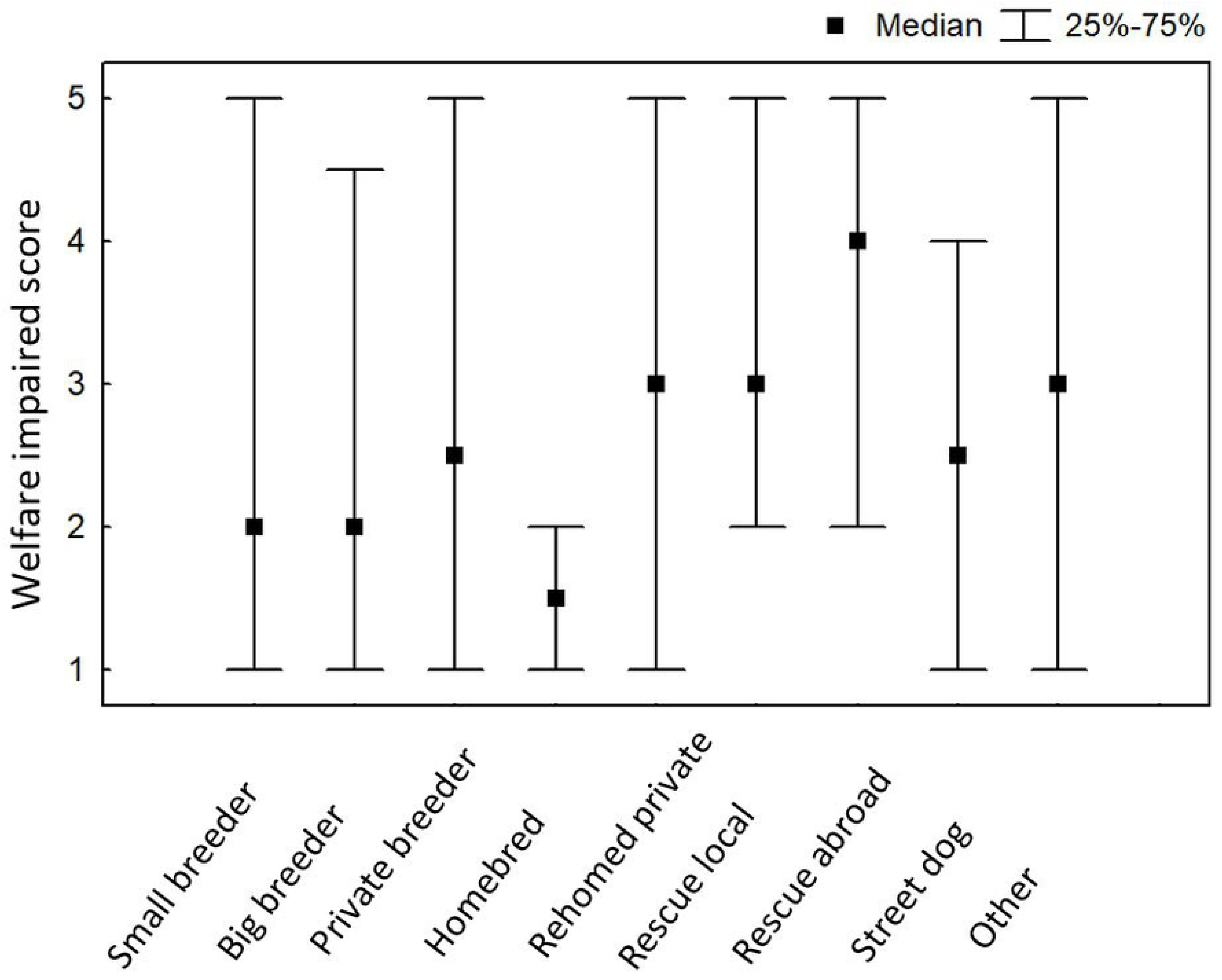
Median Welfare impaired scores and interquartile ranges for dogs from different origins.

Although there appeared to be a trend towards significance for a difference in Welfare Impaired between dogs with and without health problems (N_1_=820, N_2_=405, Manny Whitney U test, U=156252.2, p=0.093), correction for multiple testing clearly renders this result non-significant.

Results for the binomial model differed for some variables from those obtained in the univariate approach. Thus in the final “best” model, only age, breed group, health problems, and an interaction between health problems and age remained significant predictors of the occurrence of firework fears – whereas source, sex, neuter status and age at acquisition (significant predictors in the univariate analyses on severity of firework fears) were not significant, although source of dog and neuter status was still retained in the best model according to AIC (Table 3). Conversely, the model highlighted a clear effect of health problems on the occurrence of firework fears, which was not apparent in the univariate analysis, probably owing to a significant interaction between health problems and age detected in the model (Table 3, S8 Table). Thus while in the younger age groups, dogs affected by health problems had a slightly higher prevalence of firework fears, the reverse was true for the oldest age groups (Fig 3).

**Table 3.** Results of the final reduced binomial model, based on AIC, testing for the effects of health problems*age, source of dog, sex*neuter status, breed group, and age at acquisition on the occurrence of firework fears in dogs.

|  | LR Chisq | DF | p |
| --- | --- | --- | --- |
| Health problems | 16.265 | 1 | 0.00006 |
| Age | 35.380 | 10 | 0.000000003 |
| Source of dog | 12.353 | 1 | 0.262 |
| Neuter status | 2.464 | 1 | 0.117 |
| Breed group | 37.083 | 12 | 0.0002 |
| Health problems*Age | 15.807 | 1 | 0.00007 |

**Fig 3.**
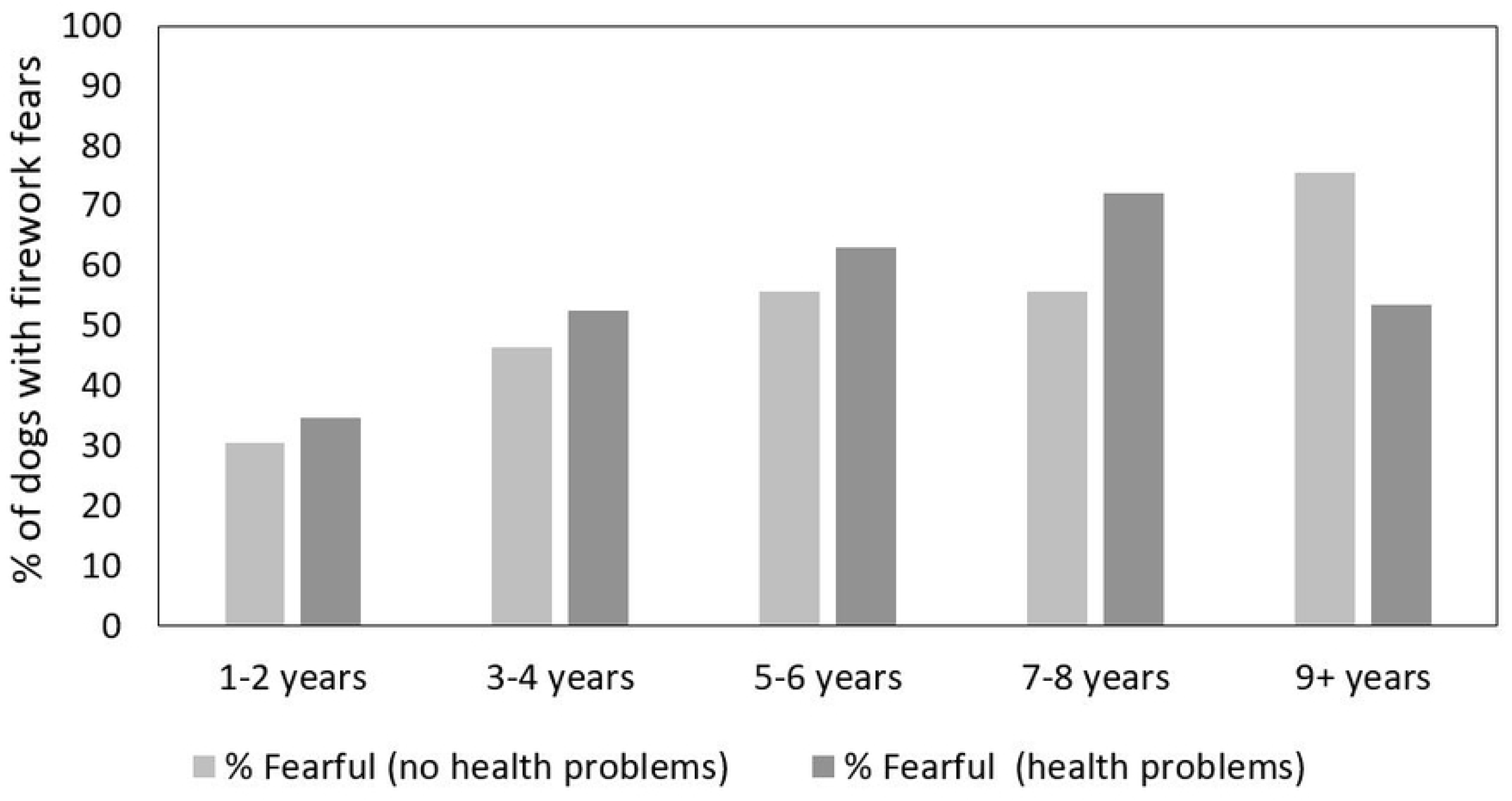
Prevalence of firework fear in dogs of different ages depending on the presence/ absence of health problems.

**Fig 4.**
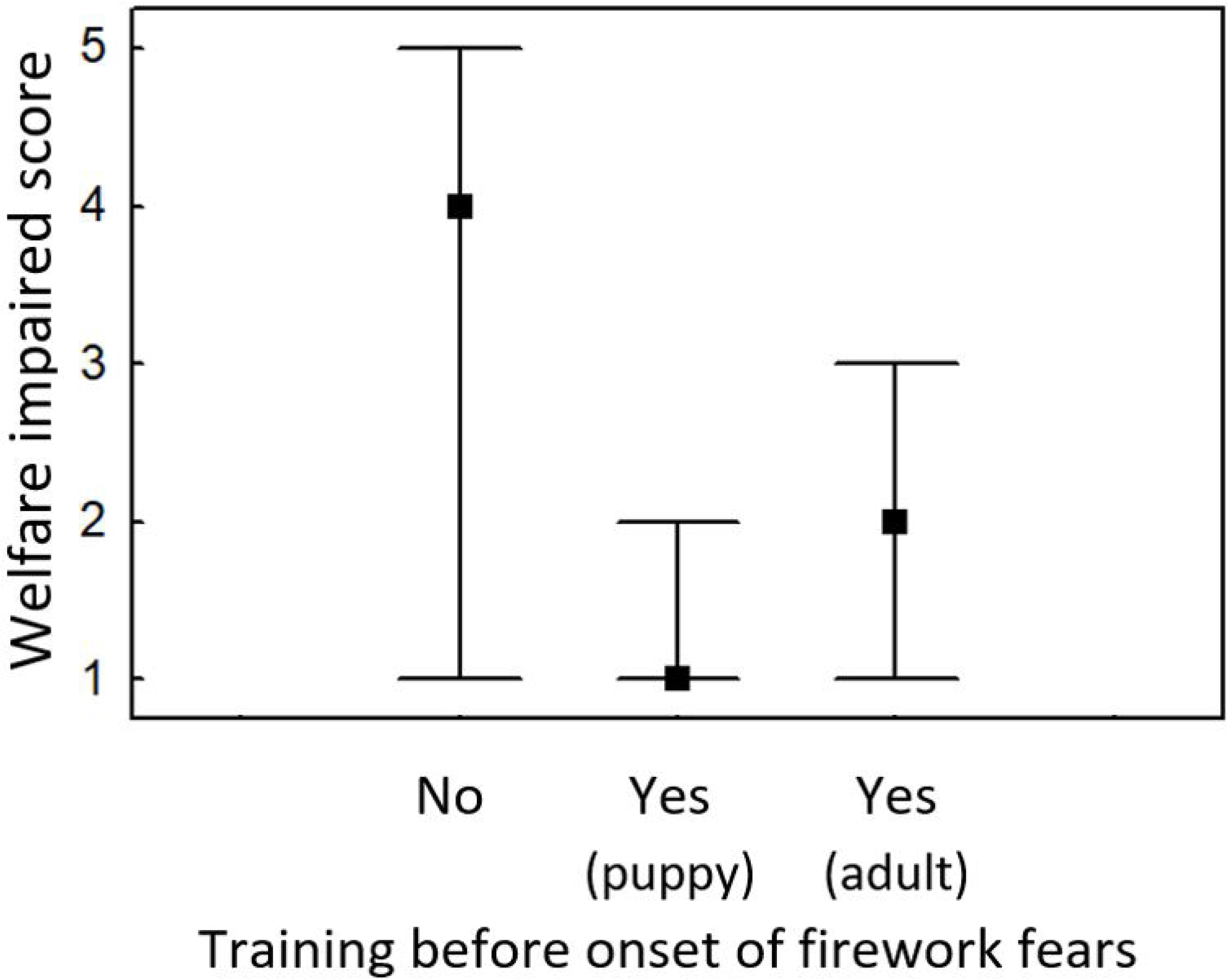
Median Welfare impaired scores and interquartile ranges for dogs whose owners performed training against firework fears with them either as puppies or adults before the onset of any noise fears versus dogs who had received no such preventative training.

**Fig 5.**
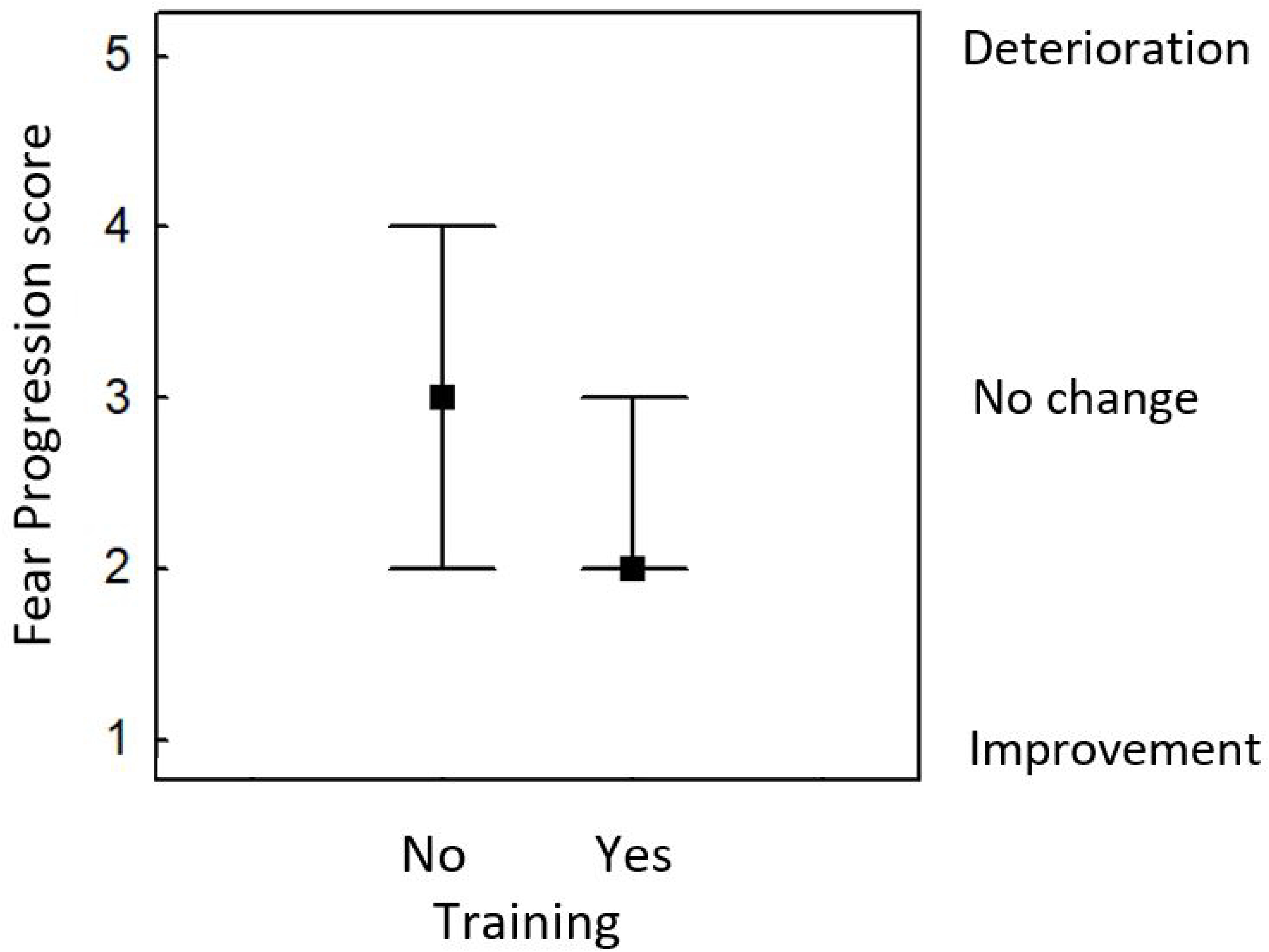
Median Fear progression scores and interquartile ranges in fearful dogs (Welfare impaired score ≥3) that had received behavioural training against firework fears vs no training.

### Relationship with other behavioural problem

#### Results of a principal components analysis with Varimax rotation on questions relating to behavioural problems

Seven principal components, explaining 83.51% of the variance, were extracted from the ‘Behaviour problems’ PCA based on biological meaningfulness (Table 4). This indicated that fear of thunder and fear gunshots loaded highly on one component (labelled “Fear of thunder/ gunshots”), while fear of other noises such as shouting and motor noise loaded on a different component (“Fear of shouting/ motor noise”). Fear and aggression towards people loaded highly on one component (“Fear/Aggression People”), with fear and aggression towards dogs loading on a different component (“Fear/ Aggression Dogs”). In contrast, the component “Resource guarding” encompassed resource guarding behaviour towards both humans and dogs. Two further components were almost entire composed of single variables, namely “Separation problems” and “Hyperactivity”, respectively. There were no cross-loadings of variables on different components, considering a cut-off point of 0.4.

**Table 4.**
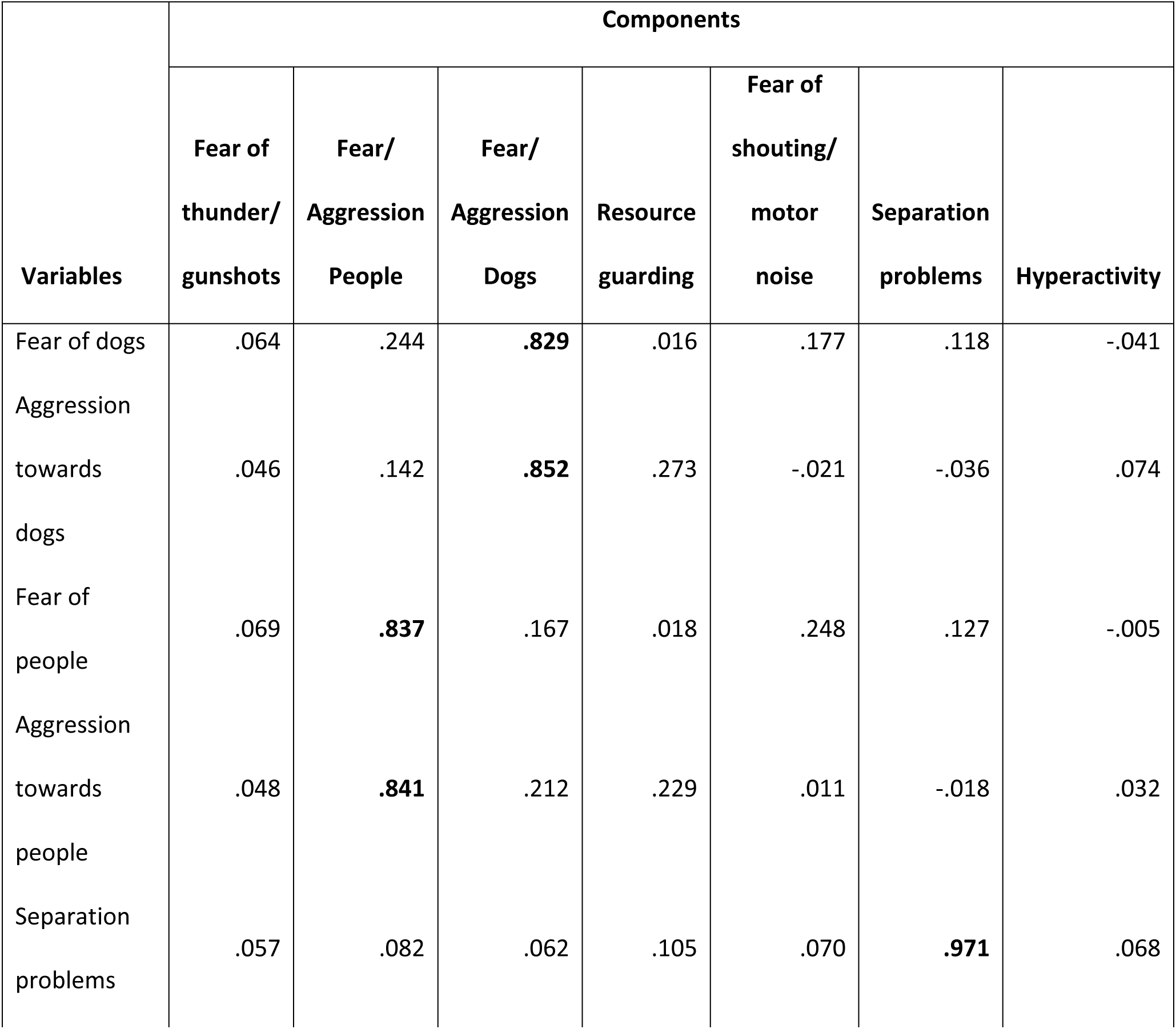
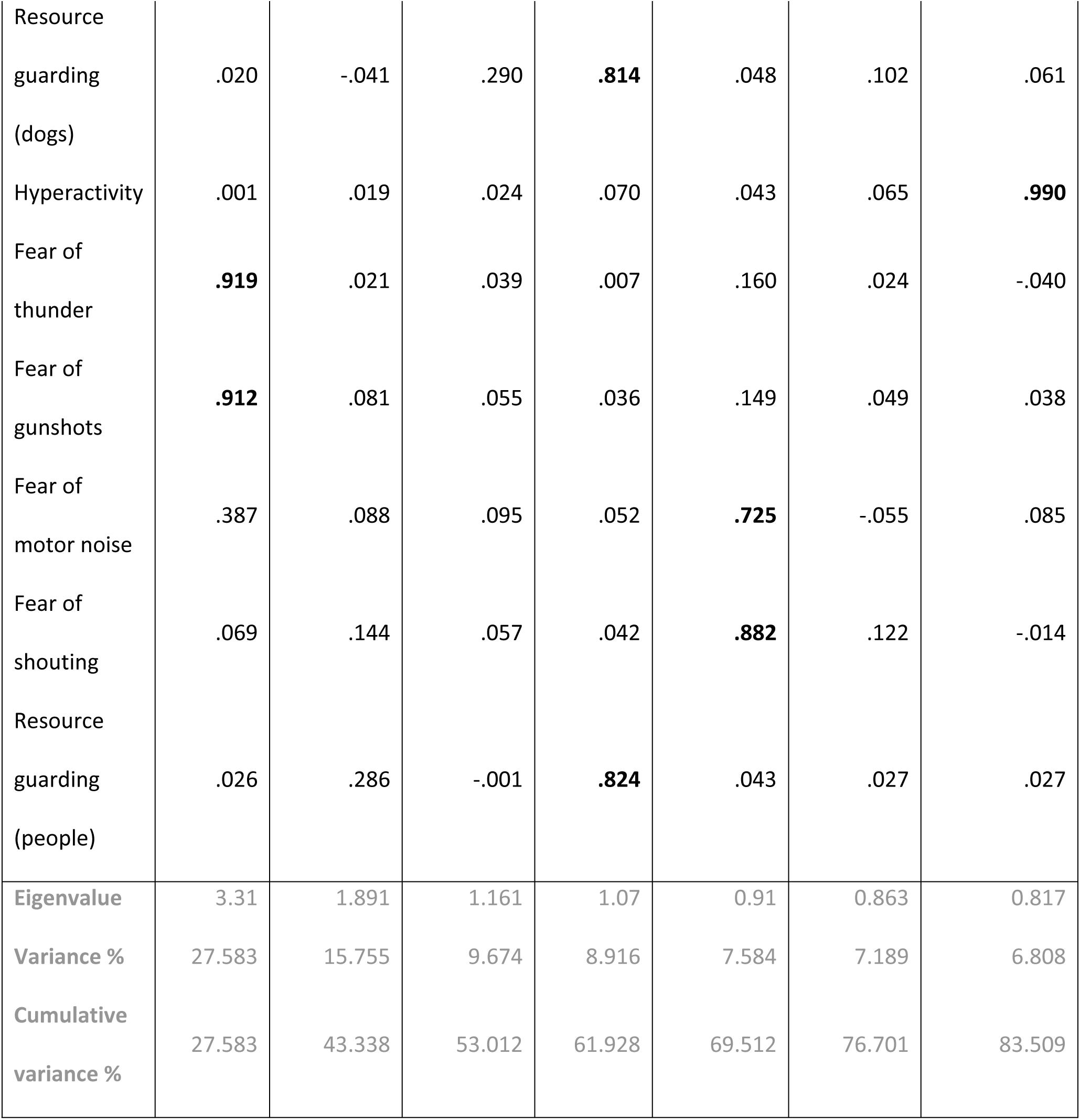
Results of a PCA on variables referring to behavioural problems other than firework fears, with Varimax rotation. Loadings >0.4 are bolded.

The Welfare Impaired score during fireworks was highly significantly positively correlated with the principal component encompassing fear of thunder and gunshots (Spearman Rho=0.719, N=1225, p<0.0000001). To a lesser extent, it was also associated with a fear of other noises such as motor noise and shouting (Spearman Rho=0.136, N=1225, p=0.000002). Although separation related problems have been suggested as a frequent co-morbidity with noise fears (29), in this sample there was no correlation with impaired welfare during fireworks (Spearman Rho=0.0003, N=1225, p=0.666). There was also no relationship with Fear/ Aggression towards people (Spearman Rho=0.027, N=1225, p=0.346), Fear/ Aggression towards dogs (Spearman Rho=0.0003, N=1225, p=0.991), Resource guarding against people and dogs (Spearman Rho=-0.018, N=1225, p=0.514), and Hyperactivity (Spearman Rho=-0.045, N=1225, p=0.119).

#### Advice sought

47.51% of owners in the total sample and 69.79% of owners of dogs with firework fears (Welfare impaired scores of 3 and above) had sought some form of advice. 36.14% of owners of fearful dogs had consulted a veterinarian, 49.76% a trainer, 46.64% the internet, 33.49% a book, 25.66% a friend, and 3.23% other sources.

#### Fear progression

As shown in Table 5, both improvement and deterioration of firework fears were frequently reported, with great improvement reported for over 10% of dogs, almost one third of dogs tending to have improved, one third of dogs with no change, just under one fifth where the fear tended to deteriorate and stark deterioration reported for 8.5%.

**Table 5.** Reported progression of firework fears in dogs affected by firework fears.

| Owners' rating of fear progression | N | % |
| --- | --- | --- |
| The fear has improved greatly | 69 | 10.88 |
| The fear tends to have improved | 180 | 28.39 |
| The fear has remained the same | 213 | 33.60 |
| The fear tends to have become worse | 118 | 18.61 |
| The fear has become much worse | 54 | 8.52 |
| **Total** | **664** | **100** |

When only dogs were included whose owners had not sought advice of any kind and did not indicate in the comments that they were behaviour specialists such as trainers or vets themselves, slightly less improvement was reported than in the full sample, but also less deterioration, with about half the dogs remaining unchanged (Table 6).

**Table 6.**
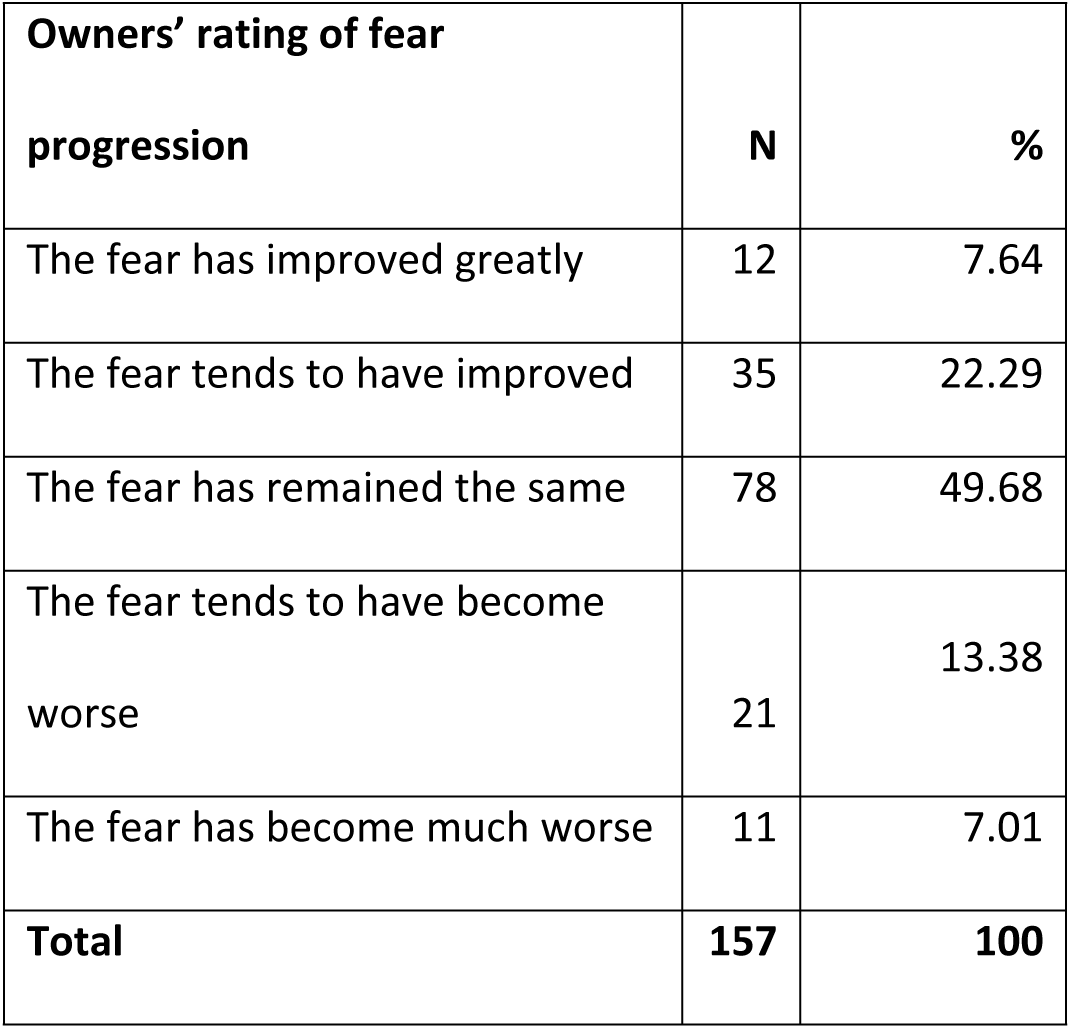
Reported progression of firework fears in affected dogs (excluding dogs with no reported firework fears), whose owners did not seek any advice to address the problem.

#### Effect of training on Welfare Impaired score

Overall, owners of 530 dogs (43.3%) had attempted some training to prevent or treat firework fears in their dogs. Regarding preventative training (before the onset of any firework fears), the owners of 228 dogs (18.8%) started to do so when their dog was a puppy and 82 (6.8%) when their dog was an adult. 74.4% did not perform any preventative training with their dogs.

The analyses demonstrated a large protective effect of training before dogs react fearfully to fireworks (Kruskal Wallis H=92.663, N=1213, p<0.0001). The median Welfare Impaired score was 1 (lowest possible score) in dogs having received training as a puppy, 2 in dogs having received training as an adult, and 4 (second highest score) in dogs with no training before the onset of any firework fears. Although the beneficial effect of training appeared to be most pronounced when training was commenced when the dog was still a puppy, post-hoc tests (two-sided significance levels with Bonferroni adjustment) indicated no significant difference in welfare scores between dogs having received training as puppies and as adults (prior to showing any fear of fireworks) (z=1.892, p=0.175). Both groups had highly significantly lower Welfare Impaired scores than dogs who received no training or whose owners commenced training only once the dogs already showed fearfulness (training as puppies: z=8.938, p<0.000001, training as adults: z=3.662, p=0.0008).

#### Effect of training on fear progression in fearful dogs

Within those dogs experiencing impaired welfare during fireworks (Welfare Impaired scores of 3 and above), the progression of firework fears was compared for dogs whose owners had attempted training against firework fears compared to no such training. The results indicated that the progression of firework fears was highly significantly more favourable in those dogs receiving training (Mann Whitney U=38853.5, N=632, p=0.00001), with a median progression score of 2 (“tended to have improved”) compared to dogs receiving no training (median progression score of 3 – “no change”).

## Discussion

In line with past studies, the results indicate that fear of fireworks is highly prevalent in the pet dog population, with 52.16% in the sample at least partly affected, and almost one third of dogs receiving the highest possible severity score. Previous studies found a prevalence of noise fears ranging from 23% to 49% (1–3). While it was explicitly stated that dogs both with and without noise fears were of interest, it is conceivable that owners with affected dogs might have a higher motivation to participate in the survey. Even if prevalence was somewhat over-estimated in our sample, the reported prevalence is consistently high also in previous studies. The majority of fearful dogs (almost 75%) had recovered by the next morning after experiencing a firework; nevertheless it took between three days to a week for full recovery in 12% of dogs, and a small proportion of dogs even took several weeks or even months to recover, with one dog’s behaviour reported to never normalise. Thus, fear of fireworks is a significant factor affecting canine welfare, both in absolute number of affected animals and duration of signs.

While no sex differences in the severity of firework fears were found, a significant effect of neutering was found in both males and females, with neutered dogs showing a greater fear of fireworks than intact dogs. This relationship appeared, however, only when analysing the effect of neutering separately as a single variable with nonparametric statistics. In contrast, when testing the effect of neutering in combination with other predictors on the presence/ absence of fireworks, neutering had no significant effect, and only the effects of breed group, age, health problems, and an interaction between health problems and age remained significant.

This may indicate that neutering per se may not actually be causative for a higher fear severity (or prevalence, as in (1,3)), but it may just be coincidental with other factors predisposing to firework fears, and this could even explain contrasting results from previous studies: (1) reported that neutered dogs (odds ratio: 1.73) were more likely to be affected by noise fears, whereas (2) found no effect of neutering. The current study used a Type 3 model, meaning that all other variables were accounted for when calculating the effect of a given predictor. The statistical methods are similar to those used by (2), who likewise included neuter status in a model with multiple predictors. On the other hand, the reported higher likelihood of noise fears in neutered dogs in (1) was based on the relative frequency of dogs fearful of noises in the neutered vs the unneutered population, and also (3) performed non-parametric tests on the effect of neuter status on behavioural signs shown during noise exposure. As such, other possible influencing factors were not taken into account. A significant effect of neutering on noise fears was found in Tiira & Lohi (30), but when only noise sensitive dogs without comorbid fears (separation related problems; fear towards strangers or in new situations) were included, this effect disappeared. Thus this effect might similarly be driven by underlying factors (such as early socialization, which became significant in the latter sample), that predisposed to a range of behavioural problems (30).

Clearly, there is still a lack of research on the behavioural effects of neutering (and at what age) in dogs – but the current study points to the importance of considering factors beyond mere correlations, with causative factors possibly only happening to coincide with neutering. Notably, the proportion of neutered individuals in our study was substantially higher in the mixed breeds (83.51%) than in the group of purebreds (or dogs from a single FCI group) at 61.04%. As can be seen in S2 table, the proportion of neutered dogs is also considerably greater in dogs originating from rescues (locally or abroad), dogs that were former street dogs, were rehomed privately, or came from another source (subsumed as “other”), compared to dogs from big breeders (5 or more litters per year), small breeders (<5 litters per year), private persons whose bitch had had a litter, or dogs that were bred and kept by their current owners. Thus, the likelihood of neutering increased with other risk factors for behavioural problems such as being of mixed breed, originating from a shelter, and an older age at acquisition. Regarding the question of effects of neutering on behavioural problems, longitudinal case control studies are needed to disentangle the effects of the surgical intervention from other potential risk factors frequently associated with neutering.

Likewise the proportion of mixed breeds compared to purebreds was much higher in dogs from rescues (locally and abroad) and street dogs compared to dogs from breeders. One large scale study (on over 15,000 dogs) has compared characteristics of mixed-breed and purebred dogs. The study found that mixed-breed dogs were more often neutered and were on average adopted at a later age than purebreds (31). Differences in owners’ demographics included that owners of mixed breeds were less educated, younger and had less experience with dogs. Mixed breeds received less training, were more likely to be kept only indoors, and as single dogs, although there was no difference in the attitude and commitment of the owners, except that time spent walking was higher for the mixed-breeds than for the purebreds (31). The incidence of problematic behaviours was significantly higher in the mixed breeds, even after controlling for the distribution of the demographic and dog keeping factors (31). The current study did not investigate whether the owners’ demographic and dog keeping characteristics differed between mixed and purebreds in the same way as in (31), but if the patterns parallel these from (31), it is possible that besides likely differences in socialisation, the lower age, education and experience with dogs of owners of mixed breeds, as well as the fact that mixed breeds received less training could contribute to the observed difference between mixed and pure breeds.

Similar to neutering, origin and age at acquisition (which had strong univariate effects) were no longer significantly associated with the occurrence of firework fears in the binomial models. Nonetheless, although the statistics point to being of mixed breed as the decisive factor, the collinearity between the predictors breed group (mixed/ pure bred), origin, age at acquisition and neutering, the data do not allow clear conclusions about the driving factor behind possible differences in fearfulness, and all of them may in fact contribute. Except for puppies adopted too early before 8 weeks of age (32,33), older ages of acquisition were found to be associated with more behaviour problems (34,35). Moreover, the early environment is particularly important in shaping dogs’ behaviour (36–38), even beyond the primary period of socialization (39,40). As also pointed out by (31), it is likely that dogs from shelters or picked up as strays, which had a much larger proportion of mixed breeds than dogs obtained from breeders, had less favourable environmental conditions during their early development. A combination of these factors may contribute to the higher incidence and severity of firework fears in mixed-breed dogs, and this might explain why univariate, but not multivariate analyses indicated significant effects of neutering, source and age at acquisition.

While mixed breeds scored the highest on the Welfare impaired score, significant differences also occurred between some breed groups. In particular, molossians, retrievers, flushing dogs and companion dogs had lower Welfare impaired scores, while herding dogs had the highest Welfare impaired scores after the group of mixed breeds. Other studies similarly found breed (group) differences, although results are not directly comparable given different breed (group) classifications (1–3).

The current study confirmed the finding by (2) that dogs that were homebred were least affected by noise fears (2). Perhaps breeders are particularly careful when socialising puppies they are intending to keep, or – having the first choice of the litter – they were most likely to keep the temperamentally most sound puppy. It is also possible that the owners who bred their own dog were especially committed, or perhaps the puppies benefitted from the older dogs in the household, with most breeders having more than one dog. Blackwell et al. (2) also suggested that breeders might be less willing to admit to the occurrence of behaviour problems in their breeding adults, or that dogs bred by their owners benefitted from remaining in the same environment they were born and socialised in.

An effect of health problems was only significant in the multivariate, but not the univariate analysis, reflecting a significant interaction between health problems and age. There was a clear increase in prevalence of firework fears with increasing age in the healthy dogs until ten years, but a decrease in dogs 11 years or older. In dogs with health problems, firework fears increased until the age of eight years and decreased in dogs nine years and older. Comparing the presence of firework fears in the two groups (dogs with/ without health problems), firework fears were moderately more common in dogs with health problems compared to healthy dogs until the age of eight years. Conversely, from the age of nine years, the incidence of firework fears was lower in dogs with health problems. Perhaps the data were less reliable for the older dogs due to the smaller sample sizes. It is possible that the proportion of dogs with health problems increased with age, but at the same time, loss of hearing, especially in the older dogs with health problems, may have attenuated some noise fears. Maybe those dogs that had health problems at an earlier age also experienced other age-related declines such as loss of hearing sooner. Thus, while health problems may contribute to firework fears at the younger ages, this effect did not appear to be strong in the current sample.

Regarding comorbidities with other behavioural problems, the current results confirm previous findings regarding the strong co-occurrence of firework fears with fears of thunder and gunshots, but a lower relationship with fears of other noises (2,3). Thus, fear of firework, gunshots and thunder does not necessarily seem to coincide with sensitivity to other types of noises. As in (2), no relationship with separation related problems was detected, although some other studies confirmed Overall et al.’s (12,29) notion that noise fears and separation related problems frequently co-occur (1,3). In my sample, no other behavioural problems were associated with firework fears, indicating that noise fears are a separate phenotype from social fears (dogs/ humans), and are also unrelated to other behaviour problems including resource guarding and hyperactivity. I did, however, not ask about fear in new situations, which was related to noise fears in (1), and was also included in a score for “Fearfulness” (together with fear toward unfamiliar people) in (3). Also, the cited studies did not use a graded score but presence or absence of noise fears. It is thus a possibility that this may to some extent account for divergent results.

In the current survey, the great majority of fearful dogs showed signs of noise fears from a very early age – 45% even developed a fear of fireworks below the age of one. Given that the age group of dogs having reached one year was the largest in the sample, data from this age group are also likely to be most reliable. The second largest group of dogs developed noise fears at the age of one year, followed by two years and three years, respectively. Very few dogs showed first signs of noise fears after the age of six years, although a new onset was reported up to the age of 12 years. Thus, the older the dog, the less likely it was to develop noise fears if it had not acquired such a fear previously.

The very early onset (median one year), as well as the observed breed differences, are suggestive of a significant genetic contribution to the development of noise fears. The median age of onset was slightly higher in Tiira et al. (3) at two years. Still, also in their sample, half of the affected dogs had developed a fear of noises in the first two years of life (3). One possible difference may lie in the form of questions. In the current study, the question was formulated as “At what age did fear of fireworks first become apparent in your dog?” This does not necessarily mean that the fear would have fully developed, and results might have been different if I had asked for manifested fears. Thus, it is likely that affected dogs do show signs from an early age, but by what Overall calls “social maturity” (at around 20 months of age in medium-sized dogs), these may have become manifest (12).

It would be expected for firework fears to increase with age, as the likelihood of encountering fear-eliciting noises inevitably increases over time (2). Furthermore, sensitisation to and generalisation of noises may occur (2). In line with this, although in most dogs, noise fears were acquired at an early age, the prevalence (1–3) and/ or average severity (4) increases with age in the population (up to a certain age), both in my sample and the cited studies. Unfortunately prevalence and severity cannot be clearly distinguished here, since if a higher number of dogs are affected by noise fears as they age (as would be expected), then the average severity score in the population would also increase, although this does not necessarily mean that individual fearful dogs show an increase in severity.

Therefore owners were also asked how their dogs’ fear of fireworks had changed in recent years. The results indicated that firework fears do not have to be a one-way road. Indeed, approximately equal numbers of respondents indicated that their dogs’ fear had improved, remained the same and deteriorated, respectively. Almost 11% even reported a great improvement, with 28% indicating some improvement. No change was noted in one third of the dogs, while the fear tended to have become worse, or had become much worse in 18% and 8% of the dogs, respectively.

The relatively high proportion of improvement is at first sight surprising. It may reflect the high proportion of owners in the current study who had sought (professional) advice – with almost half of owners of affected dogs having consulted a trainer and more than one third a veterinarian. On the other hand, even when including only dogs whose owners had not sought any advice, an improvement was reported for 30%, no change for 50%, and deterioration for 20%. This indicates that firework fears do not necessarily need to become worse over time. Also Blackwell et al. (2) report that a spontaneous recovery occurred in a small number of cases; however, for about half the cases this appeared to be due to the onset of deafness. Similarly, loss of hearing most likely explains the lower severity of firework fears in the oldest age groups in the current study.

Overall 45% of owners (and almost 70% of owners of fearful dogs) reported having sought advice (which besides trainers and vets included the internet, books or friends). This is in stark contrast to (4) where only 15.8% had sought any advice at all. Also in the sample by (2), only 29% had sought any help from vets, behaviourists, trainers, friends or other sources. This difference may reflect raised awareness of this issue over time, or may be an issue of the sampling. It cannot be ruled out that owners with a particularly high interest in dog behaviour were more likely to come across the survey invitation, which was spread in dog-related Facebook groups, while the questionnaires in the other studies were spread to a less self-selected population (contacts via veterinary practices, dog shows, agricultural or horse shows, dog walkers and other locations, (2); Auckland SPCA’s Animals Voice magazine and veterinary clinics (4)). On the other hand, both of these studies used postal surveys – requiring the owners to not only fill in the questionnaire but post it to them as well – and as such a higher effort than filing in the questionnaire online as in my study (2,4). Of interest, apart from this difference, results in the current survey regarding demographic influencing factors were remarkably similar to those obtained by Blackwell and colleagues (2).

The high proportion of owners (over 40%) who made the effort of training to address (potential) noise fears in their dogs may be another explanation for the high frequency of cases in which an improvement was noted. 25.64% had even commenced training before their dog was affected by any firework fears. This preventative training was highly successful: the median Welfare impaired score for dogs having received training as puppies was 1, meaning that the owners did not consider their dogs’ welfare to be impaired by fireworks at all. But also in adults, preventative training was useful, leading to a median Welfare impaired score of 2, compared to a score of 4 in dogs that received no preventative training.

Targeted questions indicated that owners found ad-hoc counter-conditioning (providing a high-value incentive after the occurrence of noises) and relaxation training (training dogs to relax on cue) to be the most effective training techniques for alleviating firework fears (effective in more than two thirds of cases)(41)). Thus, in a large number of dogs, prevention of noise fears by early training seems possible, and this would be a valuable piece of advice that veterinarians could give to new puppy owners when seeing the puppies for their first vaccinations, or trainers holding puppy classes. If more dog owners adopted this strategy, there would be potential to greatly reduce the incidence and/ or severity of firework fears in dogs, thus significantly improving their welfare.

## Conclusions

Older age and being a mixed breed appear to constitute the most important risk factors for firework fears in dogs. The latter might be explained by underlying differences between mixed-breed dogs and purebreds, such as in their socialisation experiences. Similarly, while severity of firework fears appears higher in neutered dogs in univariate analyses, this effect might be driven by other underlying factors, and it was no longer significant when controlling for other factors. Firework fears are highly correlated with fears of gunshots and thunder, and to a low extent with fears of other noises, but not with any other behavioural problems. Both improvement and deterioration of firework fears were frequently reported. While an early age of onset and breed differences in firework fears point to a strong genetic contribution, prevention is nonetheless possible, and training puppies as well as adult dogs to associate the noise with positive stimuli is highly effective in preventing a later development of firework fears.

## Supporting information

**S1 Table.** Relevant questions from the questionnaire survey.

**S2 Table.** Distribution of sex, neuter status and breed group (pure/ mixed) in dogs from different origins (absolute numbers).

**S3 Table.** Distribution of sex, neuter status and breed group (pure/ mixed) in dogs from different origins (percentages). Percentages ≥ 75 are shaded grey.

**S4 Table.** z-values for post-hoc tests of differences in Welfare Impaired scores in different breed groups.

**S5 Table.** p-values for post-hoc tests (adjusted for multiple testing) of differences in Welfare Impaired scores in different breed groups.

**S6 Table.** z-values for post-hoc tests comparing Welfare Impaired scores in dogs from different origins.

**S7 Table.** p-values for post-hoc tests (adjusted for multiple testing) comparing Welfare Impaired scores in dogs from different origins.

**S8 Table.** Results of a binomial model testing for the effects of Health problems*Age, source of dog, sex*neuter status, breed group, and age at acquisition on the occurrence of firework fears in dogs. Full model.

## Acknowledgements

Many thanks to all the dog owners who took the time to fill in this questionnaire, to everybody who helped to spread the survey and to Hanno Würbel for feedback on the manuscript.

